# Development and pharmacological characterization of a stable HEK293 cell model for the study of human FLT3 signaling and its pharmacological modulation

**DOI:** 10.64898/2026.10.17.759324

**Authors:** Hugo Bourki, Jean-Philippe Leyris, Sylvie Mallié, Ilana Mechaly, Pierre Sokoloff, Jean Valmier, Cyril Rivat, Hamid Moha ou Maati

**Affiliations:** Institute for Neurosciences of Montpellier, INSERM, Institut National de la Santé et de la Recherche Médicale, UMR 1298, Hôpital Saint-Eloi, Montpellier, 34000, France; Université de Montpellier, Montpellier, 34000, France; Biodol Therapeutics, Montarnaud, 34570, France

**Keywords:** Keywords : Cell line, Fms like tyrosine kinase 3 receptor, Acute Myeloid Leukemia, Chronic pain;

## Abstract

Transient transfection in adherent cell lines has been used to study the function and pharmacology of the receptor tyrosine kinase FLT3, which is involved in myeloid leukemia and pain. However, transiently transfected cells express high levels of receptors, which show ligand-independent constitutive activity. We generated a stable recombinant cell line by transfecting a plasmid construct containing the full-length Flt3 cDNA into HEK293 cells. Four individual clones, numbered 6, 10, 15 and 17 were obtained. The FLT3 receptors expressed in the four clones showed high affinity for the ligand FL, as assessed in saturation and competition binding experiments, and FLT3 activation by FL effectively initiated the recruitment of the MAPK signaling pathway, a key mediator of downstream signaling cascades for the FLT3 receptor. Clone 6 showed highest receptor activation (nearly 6-fold increase upon FL stimulation), which was selected for further pharmacological characterization. FL-induced phospho-ERK levels were inhibited by quizartinib or BDT001 with potencies like those previously reported, and showed successful phosphorylation of the FLT3 receptor assessed by TYR969. The clone N°6 of FLT3-transfected cell line therefore appears as a suitable and versatile cell line for assessing the potency of FLT3-based therapeutic candidates.

**Graphical abstract:** 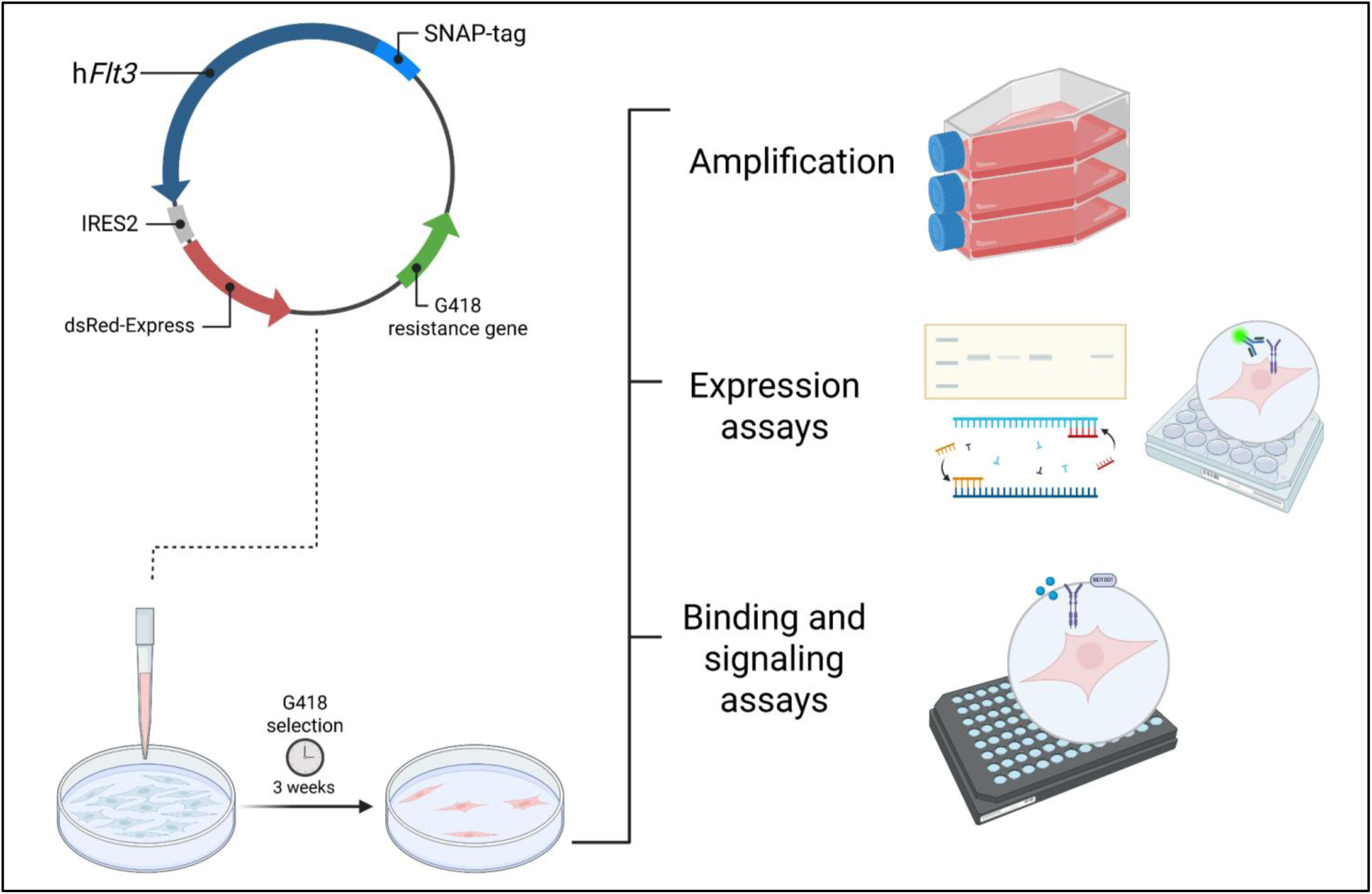

## Introduction

Tyrosine kinase receptors (RTKs) are known to play a crucial role in various aspects of cell biology. They regulate many key cellular processes such as cell survival and metabolism, cell proliferation and differentiation, as well as cell migration and control of the cell cycle (**1**). Among the different RTKs, the *Flt3* gene (Fms-like tyrosine kinase 3) encodes a type 3 RTK which plays important roles in the development and proliferation of progenitor cells of the hematopoietic lineage (**2**).

The FLT3 protein is a membrane-bound homodimer, which dimerizes upon binding of its ligand, FLT3 ligand (FL). Dimerization of the receptor triggers the autophosphorylation of its intracellular kinase domain and the initiation of several signal transduction cascades, such as the PI3K-AKT-mTOR, MAPK and JAK/STAT pathways (**2**).

Expressed in the bone marrow by cells early in hematopoietic development, FLT3 is also detected in humans in lymph nodes, thymus and spleen, however this expression mostly stops after further differentiation (**3, 4**). Mutations in FLT3 can result in constitutive receptor activation, promoting aberrant cell survival and proliferation and thereby contributing to the development of acute myeloid leukemia (AML). These mutations are detected in approximately 30% of patients with AML (**5, 6**). Consequently, FLT3 has emerged as a major prognostic marker and an important therapeutic target in AML, prompting the clinical development of several FLT3 inhibitors (**7**) We have shown that FLT3 is expressed in sensory neurons of the dorsal root ganglia, and appears to be a major actor in the initiation and maintenance of peripheral neuropathic pain (**8, 9**), as its genetical and pharmacological inhibition prevented and relieved neuropathic pain in mice models.

Human immortalized cell lines are valuable tools extensively used in cellular and molecular research, as well as in drug screening, due to their ease of cultivation. These cell lines possess cellular machinery capable of carrying out most of the post-translational folding and modifications required to produce functional and mature recombinant proteins from transfected nucleic acid sequences (**10**). While several suspension cell lines are widely used to study endogenous FLT3, such as MV4-11 for the mutated FLT3-ITD receptor and RS4-11 for the wild-type form, commercially available adherent cell lines stably expressing wild-type FLT3 are lacking. Given the interesting nature of this receptor and the pivotal roles it plays in numerous cellular processes, transient transfection is frequently employed as a default approach. However, this method is time-consuming, costly, and hampers efficient and robust pharmacological target screening due to variable transfection yields and expression levels (**11**). In addition, the lack of tight control over the membrane expression of the transfected receptor may result in non-physiological overexpression, potentially increasing ligand-independant activation, altering receptor trafficking and downstream signaling and thereby confounding pharmacological readouts. Therefore, to overcome these limitations we sought to develop a stable human embryonic kidney 293 (HEK 293) cell line expressing the FLT3 receptor. From four selected clones stably expressing FLT3, we conducted a comparative analysis with non-transfected HEK 293 cells and ultimately identified one clone as the most promising candidate. This clone was further characterized pharmacologically to confirm its suitability for downstream applications.

## Materials & Methods

### Reagents

Recombinant human FL (rh-FL) was produced in the *E. coli* Rosetta (DE3) strain (Novagen) in our laboratory using the pET15b-rhFL plasmid according to the protocol described (**12**). The rh-FL was checked for endotoxin content using the Pyrogen Recombinant Factor C endotoxin detection assay from LONZA (Walkersville MD, USA) and was found free of endotoxins. FL was labeled with a red fluorescent probe (Red-FL) by Revvity in collaboration with the Institut des Neurosciences de Montpellier (INM UR1298 INSERM). BDT001 was synthesized at Laboratoire d’Innovation Thérapeutique (UMR7200, CNRS – Université de Strasbourg, Illkirch 67400 France). Quizartinib (a.k.a AC-220), an intracellular inhibitor of Receptor Tyrosine Kinases with a high potency and a certain selectivity for FLT3 was provided by Selleckchem (#S1526). Drugs were dissolved in dimethylsulfoxide (DMSO).

### Molecular Biology

The coding sequence of human *Flt3* along with a SNAP-tag (Revvity Cisbio Bioassays) was cloned in a pIRES-dsRED2 plasmid (Vector Builders).

### Cell culture

HEK-293 cells were maintained in DMEM Glutamax (Invitrogen) supplemented with 10% heat-inactivated Foetal Bovine Serum (FBS). The culture flask was incubated at 37°C under 5% CO_2_. Stable cell lines were maintained in a medium containing a low concentration of G418 (300 µg/mL) to maintain selection pressure and limit the loss of transgene expression. Cells between passages 10 and 35 were used for experiments.

### Transfection procedures

HEK-293 cells in complete cell medium were grown in a petri dish placed at 37°C under 5% CO_2_. When reaching about 80% confluency, the cell medium was removed and replaced by a fresh cell culture medium. Then, a vector expressing hFLT3 (pIRES-SNAP-FLT3-dsREDexpress) was transfected into HEK-293 cells using JetPEI in the presence of 1 mg/ml Geneticin (G418 Sigma Aldrich). After 20–30 days of culture in G418 selection medium, individual colonies of resistant cells were isolated by using cloning cylinders (Sigma Aldrich) and grown in individual flasks containing 1 mg/mL of G418. Individual clones were then passaged two to three times before amplification and stored in liquid nitrogen in the cell medium culture containing 10% DMSO.

### Western blotting

#### Cell stimulation and protein extraction

For standard Western blot analyses, (1*10^6^) cells were collected and lysed in 150 µL of Cell Lysis Buffer (Cell Signaling Technology) for 1 h at 4 °C. Lysates were clarified by centrifugation at 10000 × g for 15 min at 4 °C, and protein concentrations were determined using a bicinchoninic acid assay (BCA; Thermo Fisher Scientific).

For FLT3 phosphorylation assays, cells were stimulated with increasing concentrations of FL for 5 min at 37 °C. Immediately after stimulation, cells were lysed in Cell Lysis Buffer supplemented with PhosSTOP phosphatase inhibitor (#4906837001; Roche) and Complete Protease Inhibitor Cocktail (#P8340; Sigma-Aldrich). Lysates were subsequently clarified and protein concentrations were determined as described above.

#### Immunoblotting

For standard protein detection, membranes were blocked for 1 h at room temperature in Tris-buffered saline containing 0.1% Tween 20 (TBST; 10 mM Tris-HCl, pH 8.0, 150 mM NaCl, and 0.1% Tween 20) supplemented with 5% non-fat dry milk. Membranes were then incubated overnight at 4 °C with anti-FLT3 (Cell Signaling Technology (#3462)) antibody diluted 1:500 in TBST containing 5% bovine serum albumin (BSA), or with anti-β-tubulin (Sigma-Aldrich (#T6199)) antibody diluted 1:2000.

For phospho-FLT3 detection, membranes were blocked for 2 h at room temperature in TBST containing 5% BSA and incubated overnight at 4 °C with anti-phospho-FLT3 (Tyr969) (Cell Signaling Technology (#3463)) antibody. Following primary antibody incubation, membranes were washed three times with TBST at room temperature and incubated with the appropriate horseradish peroxidase-conjugated secondary antibody for 1 h at room temperature for standard immunoblotting or 2 h at room temperature for phosphorylation assays. Goat anti-mouse IgG (#115-035-003; Jackson ImmunoResearch) and goat anti-rabbit IgG (#111-035-144; Jackson ImmunoResearch) were diluted 1:10000 in TBST.

#### Membrane stripping and detection of total FLT3

After phospho-FLT3 acquisition, membranes were stripped using Restore™ PLUS Western Blot Stripping Buffer (#46430; Thermo Fisher Scientific) according to the manufacturer’s instructions. Membranes were then re-blocked and incubated overnight at 4 °C with rabbit anti-FLT3 antibody. After washing, membranes were incubated for 2 h at room temperature with IRDye® 680RD donkey anti-rabbit IgG secondary antibody (#926-68073; LI-COR Biosciences).

#### Signal acquisition and quantification

Horseradish peroxidase-dependent signals were detected using SuperSignal™ West Atto Ultimate Sensitivity Substrate (Thermo Fisher Scientific) or WesternBright™ ECL HRP Substrate (#W100B and #W101B; Promega). Chemiluminescent images were acquired using a Fusion FX imaging system (Vilber). Fluorescent membranes were imaged using an Odyssey® imaging system (LI-COR Biosciences). Band intensities were quantified using Image Studio software (LI-COR Biosciences). Phosphorylated FLT3 levels were normalized to the corresponding total FLT3 levels.

#### Antibodies

The following primary antibodies were used: anti-FLT3 and anti-phospho-FLT3 (Tyr969) antibodies from Cell Signaling Technology (#3462 and #3463, respectively), and anti-β-tubulin antibody from Sigma-Aldrich (#T6199).

### Immunocytochemistry

2×10^4^ cells were seeded in 24 well plates on 14mm glass coverslips, and post-fixed in 4% paraformaldehyde after two warm Ca^2+^/Mg^2+^-free PBS (PBS^-^) washes. Cells were then blocked and permeabilized in PBS^-^ containing 10% donkey serum and 0.1% Triton x-100 during 30 min. Cells were then incubated with primary antibodies, at 4 °C overnight in PBS containing 1% donkey serum and 0.1% Triton X-100. After extensive wash in warm PBS (3 times for 10 min each) cells were incubated with appropriate secondary antibody conjugated to AlexaFluor and Hoechst (Sigma, 1 μg/ml), in the same buffer as the primary antibodies, at room temperature for 1 h and then washed (three times for 15 min each) before mounting with Mowiol. Images were acquired under a Zeiss LSM 880 Confocal microscope with ZEN software (Carl Zeiss Microscopy).

### RNA extraction and RT-qPCR

5×10^4^ cells were isolated and suspended in TRI reagent at -80 °C until RNA extraction with the TRI reagent/ chloroform technique following the protocol of the manufacturer (Sigma-Aldrich). After extraction, samples were treated with DNAse (M610A, Promega) and reverse transcribed. q-PCR reactions were made in 96 well plates with a final volume of 10 µl composed of 3 µl cDNA (final dilution 1/45), 2µL of forward and reverse primers (2.5 µM) and 5 µl of SybrGreen I master mix (Roche Diagnostics). Measures were realized with a LightCycler 480 (Roche Diagnostics). The relative amounts of amplified cDNAs were normalized with GAPDH as a stable control gene for HEK cells.

Sequences of the primer pairs used are as follows :

GAPDH : F-GATTTGGTCGTATTGGGCGC | R-TTCCCGTTCTCAGCCTTGAC

FLT3 : F-CCCAGTCAATCAGCTTTGGT | R-CCTGGCTGGTGCTTATGATT

### Covalent labeling of cells expressing the SNAP tagged-FLT3 for binding assay

After removal of the cell culture medium from a flask containing adherent cells, 10 ml of Tag-lite labelling medium containing 75 nM of SNAP-Lumi4-Tb was added to the flask and incubated for 2 h at 37°C under 5% CO_2_. The excess of SNAP-Lumi4-Tb was removed by washing each flask 4 times with 10 ml of Tag-lite labelling medium. Cells were detached, pelleted by centrifugation (5 min at 1,300 rpm) and suspended in a cell culture medium containing 10% DMSO. Then, cells were distributed at 3 million cells per vial and slowly frozen to -80°C in isopropanol and transferred in liquid nitrogen for storage. Prior to their use, the frozen cells were thawed quickly at 37°C, the medium was removed from the vials and the cells were suspended in the Tag-lite labelling medium at a cell-density of 1 million cells/ml.

### Tag-lite binding assays

When using frozen cells, binding assays were carried out immediately after cell thawing. 10,000 cells in suspension per well were used to carry out binding assays in 384-well small volume plates. All compounds used in binding assay were diluted in Tag-Lite labeling medium. Saturation binding experiments were performed in 384 wells plates by incubating the cells with increasing concentrations of Red-FL diluted in the Tag-lite labelling medium. For each concentration, nonspecific binding was determined by adding 0.1 μM of unlabeled FL diluted in the same buffer. In 384 wells plates containing 5µl of Red-FL, 5µl of unlabeled FL or Tag-lite labelling medium were added, followed by the addition of 10µl labelled cells. Plates were incubated at room temperature overnight before signal detection.

For competition assays, both Red-FL and the compounds to be tested were diluted in Tag-lite labelling medium. Cells were incubated with 0.5 nM Red-FL in the presence of increasing concentrations of compounds to be tested. In 384 wells plates containing 5µl of Red-FL, 5µl of unlabeled FL or Tag-lite labelling medium or BDT were added, followed by the addition of 10µl labelled cells. Plates were then incubated at room temperature overnight before signal detection.

HTRF signal was detected using an advanced fluorescence microplate reader (PHERAstar plus, BMG Labtech) equipped with a HTRF optic module allowing a donor excitation at 337 nm and a signal collection both at 665 nm and at 620 nm. A frequency of 40 flashes/well is selected for the laser excitation. The signal was collected both at 665 and 620 nm using the following time-resolved settings: delay 50 µs, integration time 400 µs. HTRF ratios were obtained by dividing the acceptor signal (665 nm) by the donor signal (620 nm) and multiplying this value by 10,000. The 10,000 multiplying factor is used solely for the purpose of easier data handling.

### ERK1/2 HTRF assays

HTRF experiments were done using a Phospho-ERK (Thr202/Tyr204) cellular kit (Revvity). HEK-293 clones were seeded in a 96-well plate at a concentration of 4×10^4^ cells per well following a one hour coating using poly-ornithine (Sigma-Aldrich). 24 hours later, cells were incubated 10min with FL at 37°C or 90min with antagonists prior stimulation with FL. Then, cells are immediately lysed and transferred on a 384-well plate for antibody incubation and HTRF detection, as described by the manufacturer protocol. The HTRF ratio (665/620 nm) was calculated and normalized relative to the highest concentration of FL. The sigmoidal curves were fitted to the dose–response data using non-linear regression [Log (ligand) versus HTRF ratio].

### Data analysis

K_d_ values of the fluorescent ligand Red-FL were determined from saturation curves of the specific binding (total minus non-specific binding) by using GraphPad Prism software for a one-site saturation curve.

K_i_ values of the compounds were determined from binding competition experiments according to the Cheng and Prusoff equation.

EC_50_ and IC_50_ values of the rh-FL and compounds were determined from sigmoidal dose-response curves using GraphPad Prism software with the equation :

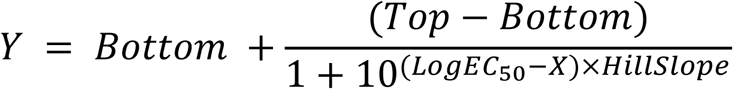

## Results

### 1 FLT3 is properly expressed in isolated HEK clones

To assess the expression of FLT3 in stably transfected HEK cells, we performed Western-blotting analysis on all isolated clones using an intracellular anti-FLT3 antibody (data not shown). Four candidates were selected based on the appropriate expected molecular weight for FLT3 fused with its SNAPtag (∼150 kDa), proper expression, and varying protein expression levels. We then blotted them again together with the appropriate untransfected and mock-transfected controls **(Fig 1A)**. The four selected clones displayed varying FLT3 expression levels compared with transiently transfected cells, whereas mock-transfected and non-transfected cells showed no detectable FLT3 expression.

**Figure 1.**
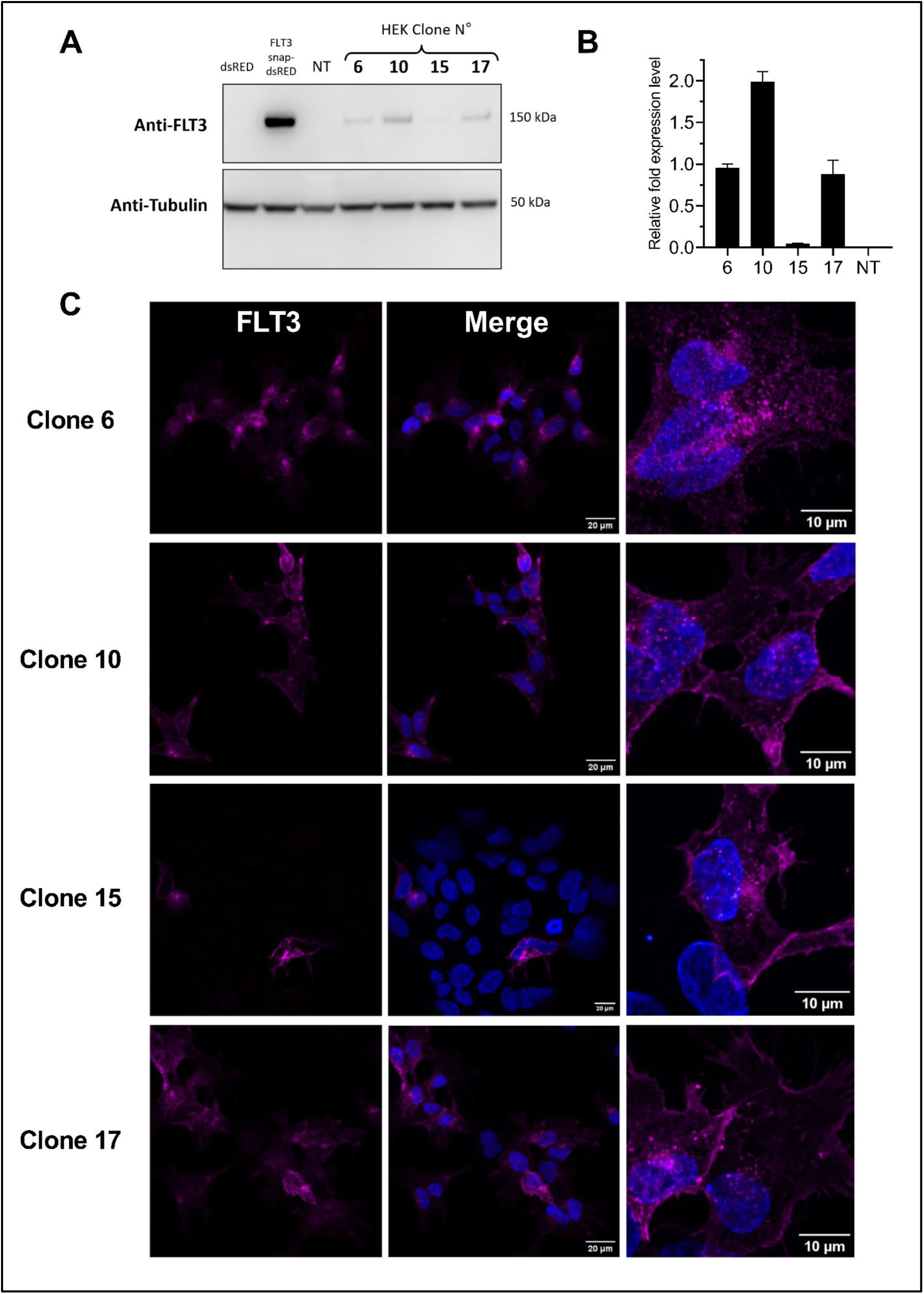
FLT3 is properly expressed in isolated HEK clones. **A:** Western blot analysis showing FLT3 expression in all stably transfected HEK clones, as bands of expected size around 150 kDa. 20µg of protein were loaded per well. Tubulin was used as a stable housekeeping gene. **B** : Relative mRNA expression levels of FLT3 by stably transfected HEK cell clones. 5×10^4^ cells were used for each clone. Gene expression levels were normalized with GAPDH. Data is represented as mean ± SD (n=2). **C** : Immunocytochemistry of HEK cell clones stably transfected by FLT3. Scale bars = 20 and 10µm.

Then, we validated and quantified the expression of *Flt3* mRNA using RT-qPCR **(Fig 1B)** with specific human FLT3 primers. Clone n°10 shows the highest relative fold expression level, followed by clones n°17, 6, and 15. *Flt3* mRNA was detected at very low levels in non-transfected HEK cells, although these values are not visible in the graph. Amplification in non-transfected cells generated the specific amplification product but only at later cycles (Ct ≈ 32). Finally, we further confirmed the expression of FLT3 in the HEK293 clones by performing immunocytochemistry **(Fig 1C)**. All four clones showed FLT3-positive staining located close to the plasma membrane. However, a proportion of cells in clone 15 lacked detectable expression, indicating heterogeneity within this population regarding the expression of FLT3.

### 2 Evaluation of the binding properties of the receptor

The affinity of Red-FL ligand for Snap-tagged FLT3 was first determined by binding saturation experiments performed on each HEK FLT3 clone. Increasing concentrations of Red-FL increased the HTRF signal until a plateau, which indicates saturation of FLT3 binding. The calculated dissociation constants (Kd) were 0.10, 0.19, 0.06 and 0.35 nM for clones n°6, 10, 15 and 17, respectively (**Fig 2A**). Non-specific binding varied linearly with increasing Red-FL concentrations. Subsequently, we compared the ligand-binding properties of FLT3 across the four selected HEK293 clones by determining the ability of FL to displace Red-FL from FLT3 over a wide concentration range. Increasing concentrations of FL resulted in a dose-dependent reduction in specific binding of Red-FL, yielding IC50 values ranging from 1 to 3 nM (Fig 2B). These results show that FL binds to FLT3 with high affinity, with closely comparable IC₅₀ values across the four FLT3-expressing HEK293 clones.

**Figure 2.**
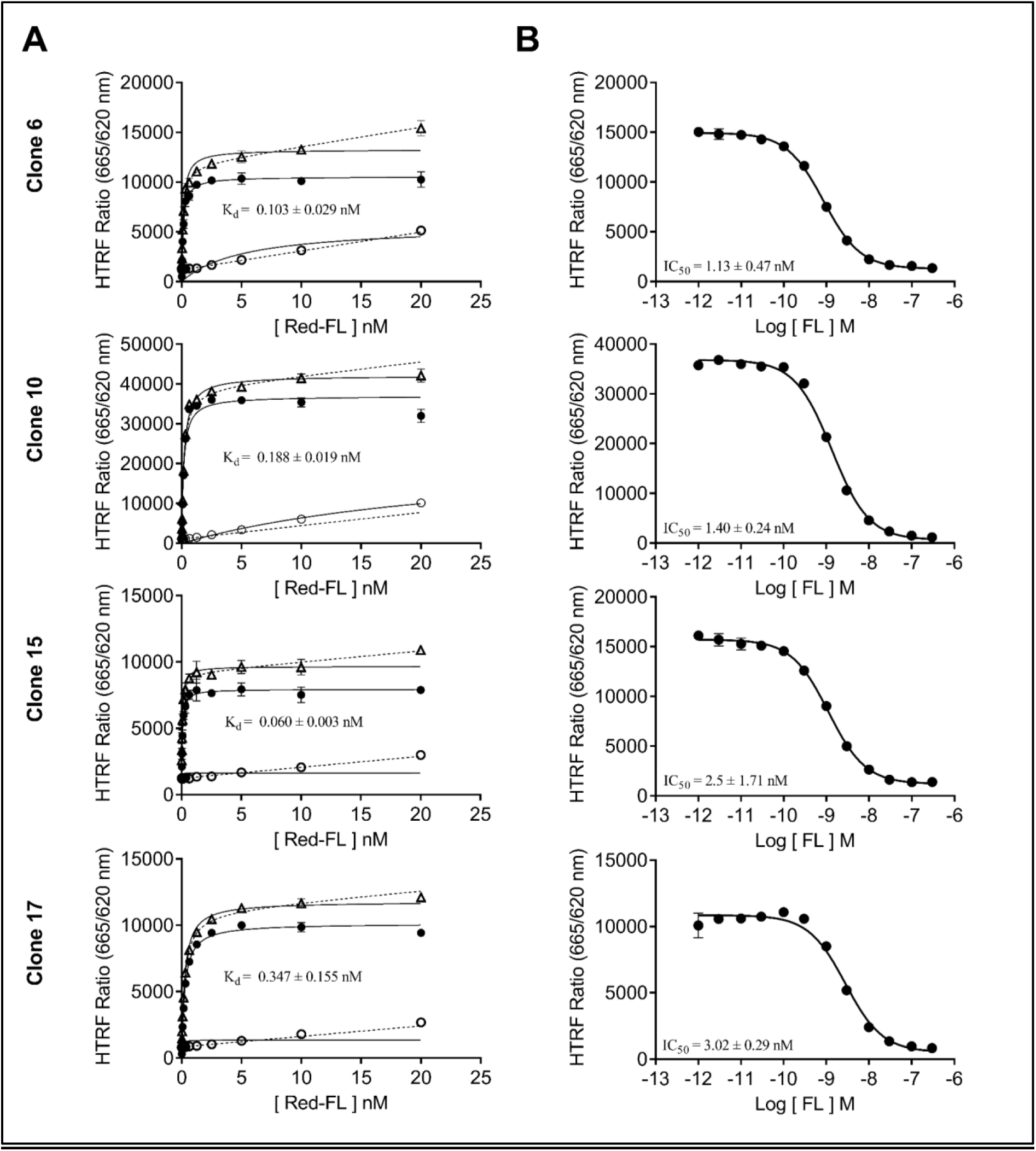
FLT3 binds to FL with high affinity and specificity. A : Saturation binding assays on Snap-FLT3. The specific binding signal (filled circle) was calculated by subtracting the nonspecific binding signal determined in the presence of 0.1 µM unlabeled FL (open circle) from the total signal (open triangle). Plotted values are representative of one experiment. K_d_ values are mean ± SEM of two experiments (n=2), each performed in triplicate. | B : Competition binding assays on Snap-FLT3. Plotted values are representative of one experiment. IC_50_ values are mean ± SEM of two experiments each performed in triplicate (n=2) and were calculated using the K_d_of each corresponding saturation binding experiment, according to the Cheng and Prusoff equation.

### 3 Evaluation of the signaling properties of the receptor

Next, we examined whether binding of FL and subsequent receptor activation effectively initiated the recruitment of the MAPK signaling pathway, one of the key mediators of downstream signaling cascades for the FLT3 receptor. We used Homogeneous Time Resolved Fluorescence assay (HTRF) to assess intracellular ERK1/2 phosphorylation levels. Stimulation with FL led to a dose-dependent increase in the HTRF signal, indicating phosphorylation of ERK1/2 (**Fig 3A**) for all HEK clones, with an EC_50_ varying between 0.3 and 1.5 nM. The extent of FL-induced activation of the ERK1/2 pathway, expressed as the top to bottom response ratio varied amongst clones: 5.89, 2.49, 1.10 and 4.81, for clones 6, 10, 15 and 17, respectively. The specificity of the response was confirmed by the low amount of ERK phosphorylation in cells that are non-stimulated.

**Figure 3.**
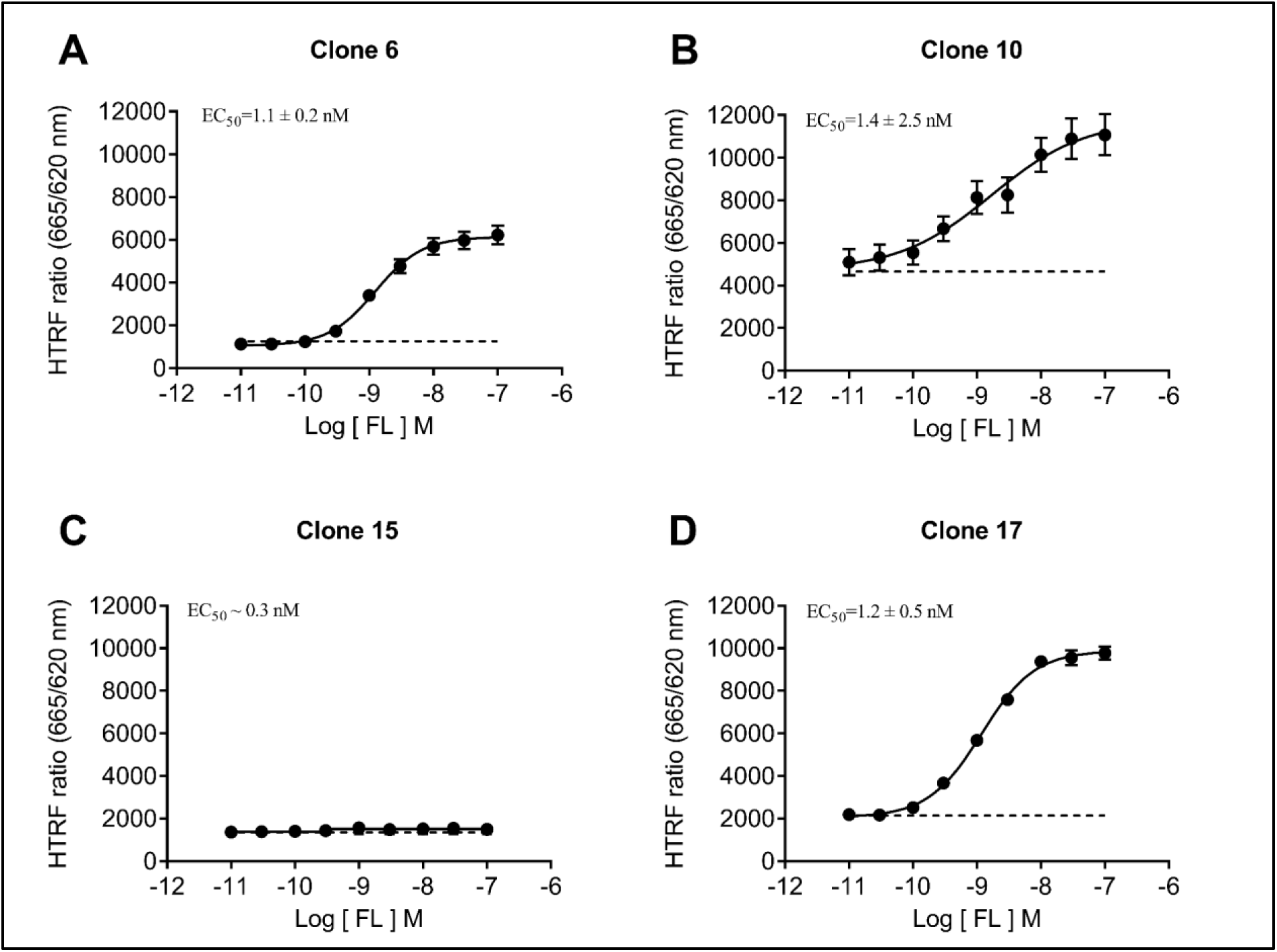
ERK1/2 signaling using HTRF assays. A : Phospho-ERK1/2 assays under FL stimulation performed on each clone. The percentage of normalized Phospho-ERK levels were calculated relative to the highest HTRF ratio observed in each experiment. Plotted values are representative of one experiment. EC50 values are mean ± SEM of three independent experiments (n=3), each performed in triplicate. Dashed lines represent baseline HTRF levels.

**Figure 4.**
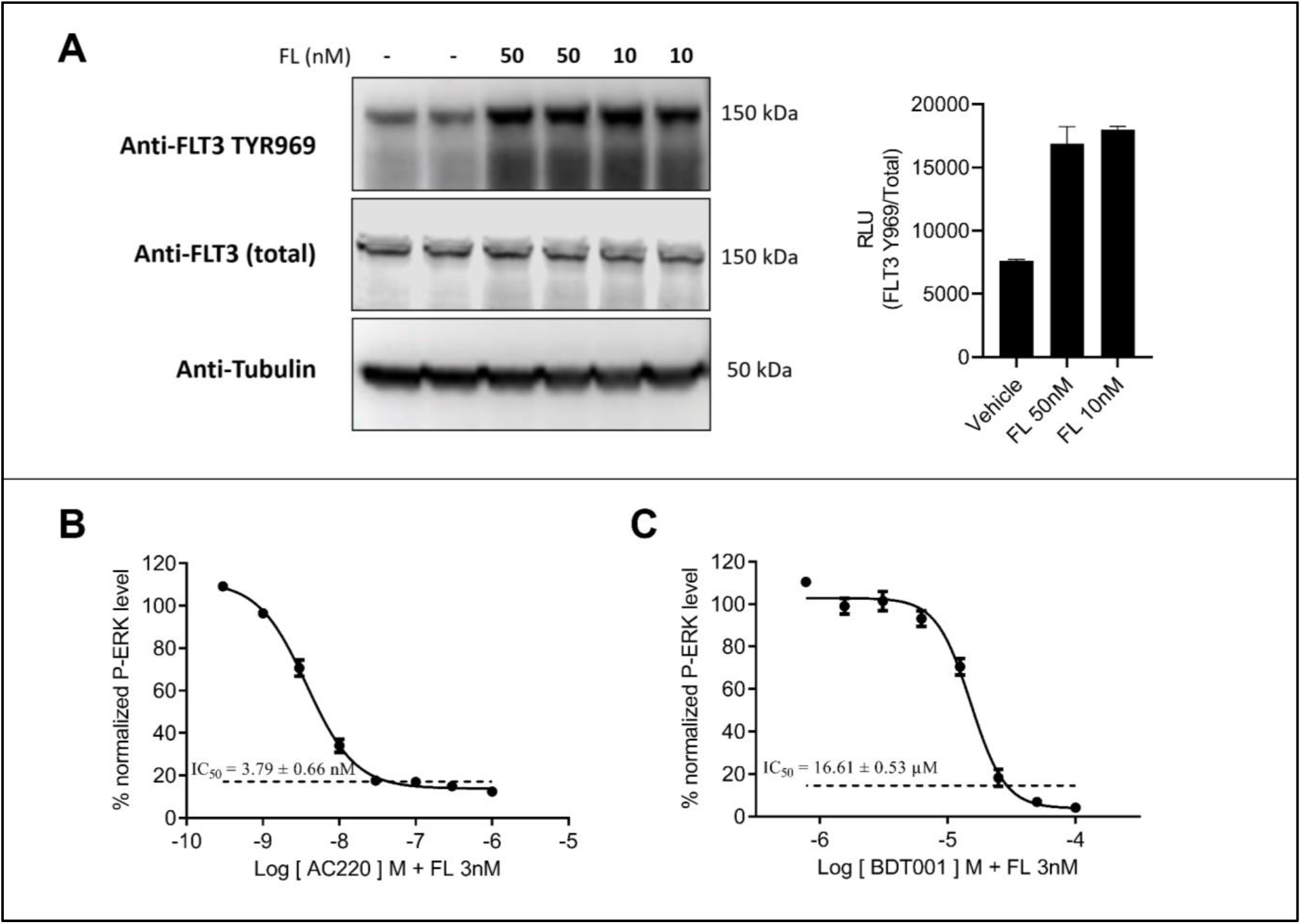
FLT3-expressing HEK clone n°6 conserves pharmacological properties of the native receptor. **A :** Western blot analysis showing phospho-FLT3 expression in HEK clone n°6 (*left*) along with the phospho-FLT3/total FLT3 quantification ratio (*right*) (n=2 replicates from the same sample). 20µg of protein were loaded per well. Tubulin was used as a stable housekeeping gene **B and C :** Phospho-ERK1/2 assays under FL stimulation and AC220 or BDT001 inhibition performed on each clone. The percentage of normalized Phospho-ERK levels were calculated relative to the HTRF ratio for FL 3nM. IC50 values are mean ± SEM of three independent experiments for AC220 (n=3) and two for BDT001 (n=2) each performed in triplicate. Dashed lines represent baseline HTRF levels

Taken together, these results show that our HEK clones expressing FLT3 are able to effectively recruit the MAPKinase intracellular signaling pathway confirming the functional activity of the expressed receptors. When comparing the different clones for expression and functional activity, clone 6 was the most responsive to FL stimulation for the recruitment of the ERK1/2 MAPK signaling pathway (near 6-fold increase) and showed robust membrane expression on immunocytochemistry. Therefore, clone N°6 was selected for further pharmacological assays.

### 4 Pharmacological validation of Clone n°6

We first assessed FL-induced phosphorylation of FLT3 by Western blotting using a phospho-specific antibody targeting Tyr969. Treatment with 10 or 50 nM FL produced an approximately threefold increase in the phospho-FLT3/total FLT3 ratio, indicating robust FL-dependent phosphorylation of the FLT3 receptor.

We next assessed the inhibition of FL stimulation by quizartinib, an intracellular kinase inhibitor of FLT3, displaying some selectivity for this receptor (**6**) or by BDT001, a negative allosteric modulator of FLT3 (**2**). Both compounds dose-dependently inhibited the HTRF signal, indicating decreased levels of ERK1/2 phosphorylation, with IC_50_ values of 3.8nM and 16.6µM for quizartinib and BDT001 respectively. Thus, FLT3 expressed by clone n°6 retained its pharmacological sensitivity to inhibition by quizartinib and BDT001. Together, these findings identify clone 6 as a suitable cellular model for assessing the pharmacological inhibition of FLT3 activity by compounds with distinct mechanisms of action.

## Discussion

Using non-targeted gene delivery, we generated a panel of HEK clones expressing the tyrosine kinase receptor FLT3.

First, we confirmed the quantitative expression of the receptor both at the mRNA and protein level using RT-qPCR and Western blotting for the 4 selected clones, showing variable amounts of receptor expression among the clones. Variability in expression levels was expected due to the non-targeted nature of transgene integration, as transcriptional activity can differ depending on the genomic insertion site and its local regulatory environment (**13**). Immunocytochemical analysis further confirmed FLT3 protein expression in the selected clones and revealed a predominantly extra-nuclear distribution, characterized by diffuse and punctate cytoplasmic staining, including peri-nuclear immunoreactivity, along with some signal extending toward the cell periphery and cellular processes. Such a distribution is compatible with the presence of both intracellular and membrane-associated receptor pools. This is consistent with the biology of receptor tyrosine kinases, which undergo continuous intracellular trafficking between the plasma membrane and intracellular compartments, including endosomes, as part of receptor maturation, internalization, recycling and degradation (**14**, **15**). Intracellular signals could therefore reflect receptors undergoing post-translational trafficking or post-endocytosis sorting. However, in the absence of co-localization with plasma membrane or organelle-specific markers, the precise subcellular compartments containing FLT3 cannot be determined from our experiments. Of note, clone n°15 appears to consist of a heterogeneous population of FLT3-positive and FLT3-negative cells, both resistant to G418, as shown by immunocytochemistry staining. This heterogeneity could be due to a subset of cells losing FLT3 expression over time, with FLT3-negative cells gaining a selective advantage by avoiding the metabolic burden of protein production. This would allow them to proliferate more rapidly within the population, and is consistent with our use of non-targeted gene delivery. Alternatively, contamination from a neighboring clone or non-adherent cells during isolation with cloning cylinders could also account for the observed heterogeneity.

While proper expression of FLT3 was confirmed, it was also necessary to determine whether the recombinant receptor retained its ligand-binding properties. Saturation and competition assays demonstrated sub-nanomolar affinity of SNAP-FLT3 for Red-FL, supporting the preservation of high-affinity ligand recognition. Nevertheless, the use of a SNAP-tagged recombinant receptor introduces several limitations, as fusion tags and fluorescent-tagged ligands may potentially alter receptor or ligand properties. Nevertheless, K_d_ values for FL obtained using our cell lines are consistent with described values using cell lines that express FLT3 endogenously (**16**). The preservation of labeled-ligand binding, together with competition by unlabeled ligands, argues against a major functional alteration of the receptor. At the functional level, we measured effective downstream phosphorylation of the ERK1/2 signaling pathway after FL stimulation. All clones showed variable ERK1/2 phosphorylation baseline levels and FL-dependant activation range, with an EC_50_ consistent with previously reported values (**17**). Clone 15 showed the lowest FL-dependent ERK1/2 phosphorylation ratio, which might be explained by the presence of FLT3 negative cells in the culture. Since signal detection is averaged across the entire well, a small population of FLT3-positive cells may result in insufficient ERK1/2 activation, making it difficult to distinguish from baseline levels containing FLT3-negative cells. Moreover, other signaling pathways, such as PI3K-AKT-mTOR and JAK/STAT, were not investigated. Therefore, we cannot exclude the possibility that individual clones exhibit pathway-selective signaling or differential signaling bias across downstream pathways.

The optimal clone of interest suited for drug screening and other downstream applications is one with good receptor binding kinetics, consistent and efficient recruitment of its intracellular signaling pathways, and a balanced expression of the receptor at the plasma membrane. The latter reason is especially important considering the fact that type 3 tyrosine kinase receptors, when overexpressed at the membrane, can promote ligand-independent activation, consistent with the idea that membrane overexpression increases the probability of spontaneous receptor proximity/dimerization and autophosphorylation (**18, 19**). Among the FLT3-positive clones isolated, Clone n°6, which displayed the highest extent of stimulation by FL, was selected as a cell line to further characterize FLT3 inhibition. While BDT001 acts as a non-competitive extracellular negative allosteric modulator **[8]**, Quizartinib binds to the intracellular activation loop and inhibits kinase activity **[20]**. Despite their different modes of action, both Quizartinib and BDT001 inhibited FL-stimulated ERK phosphorylation, with potencies similar to those previously reported **[8, 20]**. The distinction in their mechanisms of inhibition helps to attest the receptor’s structural integrity and functionality. Finally, ligand-dependent phosphorylation of FLT3 at TYR969 was validated by Western blotting using a phospho-specific antibody. However, phosphorylation at additional tyrosine residues sites was not assessed. Although our results align with initial expectations, some unanticipated outcomes deserve further discussion. Upon confocal microscopy analysis, the four selected clones did not exhibit red fluorescence, indicating a lack of ds-RED expression. This could be explained by our selection method, where clones were selected based on FLT3 expression through Western blotting, which may have included the potential disruption of IRES or dsRed sequences during genomic integration. Finally, *Flt3* is not typically expressed in HEK cells, but was unexpectedly detected at low levels in untransfected cells using qPCR, possibly due to ectopic expression.

To conclude, we developed in the present study a FLT3-expressing stable HEK cell-line and characterized it across passages ranging from 10 to 35 to ensure its functionality and stability. We confirmed the proper expression of the receptor both at the mRNA and protein level using RT-qPCR, Western blotting and Immunocytochemistry, we assessed appropriate binding kinetics to its ligand using saturation and competition binding experiments, assessed ligand-dependent phosphorylation of one of its tyrosine residues, and measured effective recruitment of its main intracellular signaling pathway as well as its inhibition using two different antagonists. This cell-line can be used as a tool to study the FLT3 receptor in various physiological and pathological contexts. The FLT3 construct, engineered with SNAP and Flag tags within an IRES-dsRed vector, provides a versatile platform to study the receptor’s multifaceted roles. This design enables diverse applications, such as tracking the expression of the receptor and investigating receptor-ligand interactions. This reflects the receptor’s broad relevance in many research ^areas such as^ oncology, hematopoiesis, and drug development.

## Author Contributions

**Conceptualization,** H.M., C.R. and P.S. ; **Methodology,** H.M., J.P.L., S.M., I.M. ; **Validation,** J.P.L, H.M., C.R. ; **Formal Analysis**, H.B., J.P.L., H.M. ; **Investigation,** H.B., J.P.L., H.M. ; **Data Curation,** H.B., J.P.L., P.S. ; **Writing – Original Draft Preparation,** H.B. ; **Writing – Review & Editing,** H.B., J.P.L., P.S., J.V. and C.R. ; **Supervision,** H.M. and C.R ; **Funding Acquisition,** J.V., C.R. and P.S.

All authors have read and agreed to the published version of the manuscript.

## Acknowledgments

We acknowledge the imaging facility MRI, member of the national infrastructure France-BioImaging (https://ror.org/01y7vt929) supported by the French National Research Agency (ANR-24-INBS-0005 FBI BIOGEN)

